# Convergent evolution of gene expression in two high-toothed stickleback populations

**DOI:** 10.1101/259499

**Authors:** James C. Hart, Nicholas A. Ellis, Michael B. Eisen, Craig T. Milller

**Affiliations:** Department of Molecular and Cell Biology, University of California-Berkeley, CA, USA; Howard Hughes Medical Institute, University of California, Berkeley, CA, USA

## Abstract

Changes in developmental gene regulatory networks enable evolved changes in morphology. These changes can be in *cis* regulatory elements that act in an allele-specific manner, or changes to the overall *trans* regulatory environment that interacts with *cis* regulatory sequences. Here we address several questions about the evolution of gene expression accompanying a convergently evolved constructive morphological trait, increases in tooth number in two independently derived freshwater populations of threespine stickleback fish (*Gasterosteus aculeatus*). Are convergently evolved *cis* and/or *trans* changes in gene expression associated with convergently evolved morphological evolution? Do *cis* or *trans* regulatory changes contribute more to the evolutionary gain of a morphological trait? Transcriptome data from dental tissue of ancestral low-toothed and two independently derived high-toothed stickleback populations revealed significantly shared gene expression changes that have convergently evolved in the two high-toothed populations. Comparing *cis* and *trans* regulatory changes using phased gene expression data from F1 hybrids, we found that *trans* regulatory changes were predominant and more likely to be shared among both high-toothed populations. In contrast, while *cis* regulatory changes have evolved in both high-toothed populations, overall these changes were distinct and not shared among high-toothed populations. Together these data suggest that a convergently evolved trait can occur through genetically distinct regulatory changes that converge on similar *trans* regulatory environments.

**Author Summary:** Convergent evolution, where a similar trait evolves in different lineages, provides an opportunity to study the repeatability of evolution. Convergent morphological evolution has been well studied at multiple evolutionary time scales ranging from ancient, to recent, such as the gain in tooth number in freshwater stickleback fish. However, much less is known about the accompanying evolved changes in gene regulation during convergent evolution. Here we compared evolved changes in gene expression in dental tissue of ancestral low-toothed marine fish to fish from two independently derived high-toothed freshwater populations. We also partitioned gene expression changes into those affecting a gene’s regulatory elements (*cis*), and those affecting the overall regulatory environment (*trans*). Both freshwater populations have evolved similar gene expression changes, including a gain of expression of putative dental genes. These similar gene expression changes are due mainly to shared changes to the *trans* regulatory environment, while the *cis* changes are largely population specific. Thus, during convergent evolution, overall similar and perhaps predictable transcriptome changes can evolve despite largely different underlying genetic bases.

## Introduction

Development is controlled by a complex series of interlocking gene regulatory networks. Much of this regulation occurs at the level of transcription initiation, where *trans* acting factors bind to *cis* regulatory elements to control their target gene’s expression [1,2]. Evolved changes in an organism’s morphology are the result of changes in this developmental regulatory landscape. It has been proposed that the genetic bases of many of these evolved changes are mutations within the *cis*-regulatory elements of genes [3–5]. Indeed, recent work in evolutionary genetics suggests the molecular bases of a diverse array of traits from *Drosophila* wing spots [6] to mouse pigmentation [7] to stickleback armored plate number [8,9] and size [10] are changes in the activity of *cis*-regulatory elements.

Evolved changes in gene expression can be divided into two broad regulatory classes. *Cis* regulatory changes occur within the proximal promoter [11], distal enhancer [12], or the gene body itself [13]. *Trans* regulatory changes modify the overall regulatory environment [14,15], but are genetically unlinked to the expression change. The total evolved gene expression differences can be partitioned into changes in *cis* and *trans* by quantifying expression differences between two populations and also testing for expression differences between alleles in F1 hybrids between the two populations [16]. Several studies have attempted to characterize evolved *cis* and *trans*-regulatory changes at a transcriptome-wide level [17–21]. Though the relative contribution of *cis* and *trans* regulatory changes varies extensively among studies, *cis* changes have been found to dominate [17,18,21] or at least be approximately equivalent [19,20] to *trans* changes [22]. Additionally, compensatory changes (*cis* and *trans* changes in opposing directions) have been found to be enriched over neutral models [17,18], showing evidence for selection for stable gene expression levels. However, none of these studies examined contribution of *cis* and *trans* gene expression changes during convergent morphological evolution.

Populations evolve new traits following a shift to a novel environment, due to a mixture of drift and selection. Truly adaptive traits can often be repeatedly observed in multiple populations following a similar ecological shift. Threespine sticklebacks are an excellent system for the study of evolved changes in phenotypes, including gene expression [23–27]. Marine sticklebacks have repeatedly colonized freshwater lakes and streams along the coasts of the Northern hemisphere [28]. Each of these freshwater populations has independently adapted to its new environment; however, several morphological changes, including a loss in armored plates and a gain in tooth number, are shared among multiple newly derived populations [29,30]. The repeated evolution of lateral plate loss is due to repeated selection of a standing variant regulatory allele of the *Eda* gene within marine populations [8,9] and genome sequencing studies found over a hundred other shared standing variant alleles present in geographically diverse freshwater populations [31]. These studies suggest the genetic basis of freshwater adaptation might typically involve repeated reuse of the same standing variants to evolve the same adaptive freshwater phenotype.

However, more recent evidence has shown that similar traits have also evolved through different genetic means in freshwater stickleback populations. A recent study which mapped the genetic basis of a gain in pharyngeal tooth number in two independently derived freshwater populations showed a largely non-overlapping genetic architecture [30]. Another study using three different independently derived benthic (adapted to the bottom of a lake) populations showed that, even when adapting to geographically and ecologically similar environments, the genetic architecture of evolved traits is a mix of shared and unique changes [32]. Even in cases where the same gene is targeted by evolution in multiple populations (the loss of *Pitx1* expression resulting in a reduction in pelvic spines), the individual mutations are often independently derived [33,34]. All of these genomic scale studies have looked at the genetic control of morphological changes, while the extent and nature of genome-wide gene expression changes has been less studied. It remains an open question as to whether similar gene expression patterns evolve during the convergent evolution of morphology, and if so, to what extent those potential shared gene expression changes are due to shared *cis* or *trans* changes.

Teeth belong to a class of vertebrate epithelial appendages (including mammalian hair) that develop from placodes, and have long served as a model system for studying organogenesis and epithelial-mesenchymal interactions in vertebrates [35]. Odontogenesis is initiated and controlled by complex interactions between epithelial and mesenchymal cell layers, and involves several deeply conserved signaling pathways [36–38]. Sticklebacks retain the ancestral jawed vertebrate condition of polyphyodonty, or continuous tooth replacement, and offer an emergent model system for studying tooth replacement. Previous work has supported the hypothesis that two independently derived freshwater stickleback populations have evolved an increase in tooth replacement rate, potentially mediated through differential odontogenic stem cell dynamics [30] (Cleves et al 2018 under review, see Supplementary Data File 2). Recent studies have found teeth and taste bud development to be linked, with one study supporting a model where teeth and taste buds are copatterned from a shared oral epithelial source [39], and another study supporting a model where teeth and taste buds share a common progenitor stem cell pool [40].

We sought to examine the evolution of the regulatory landscape controlling stickleback tooth development and replacement. Using high-throughput RNA sequencing (RNA-seq), we found that two independently derived high-toothed freshwater populations display highly convergent gene expression changes, especially in orthologs of known tooth-expressed genes in other vertebrates, likely reflecting the convergently evolved tooth gain phenotype and the deep homology of teeth across all jawed vertebrates. We also quantitatively partitioned these evolved gene expression changes into *cis* and *trans* regulatory changes [16,19] in both populations at a transcriptome-wide level using RNA-seq on F1 marine-freshwater hybrids. We found that *trans* regulatory changes predominate evolved changes in gene expression in dental tissue. Additionally, we found that the *trans* regulatory changes are more likely to be shared between the freshwater populations than the *cis* regulatory changes. Thus, similar downstream transcription networks controlling tooth development and replacement have convergently evolved largely through different upstream genetic regulatory changes.

## Results

### Convergent evolution of tooth gain in two freshwater populations

To further test whether multiple freshwater populations have evolved increases in tooth number compared to multiple ancestral marine populations [30,41], we quantified total ventral pharyngeal tooth number of lab reared sticklebacks from four distinct populations: (1) a marine population from the Little Campbell river (LITC) in British Columbia, Canada, (2) a second marine population from Rabbit Slough (RABS) in Alaska, (3) a benthic freshwater population from Paxton Lake (PAXB) in British Columbia, Canada, USA, and (4) a second freshwater population from Cerrito Creek (CERC) in California, USA (Fig 1A, 1B). Freshwater fish from both populations had more pharyngeal teeth than marine fish at this 35-50mm standard length (SL) stage, consistent with previous findings [30,41] of increases in tooth number in freshwater sticklebacks (Fig 1B, 1C, Table S1).

**Fig 1.**
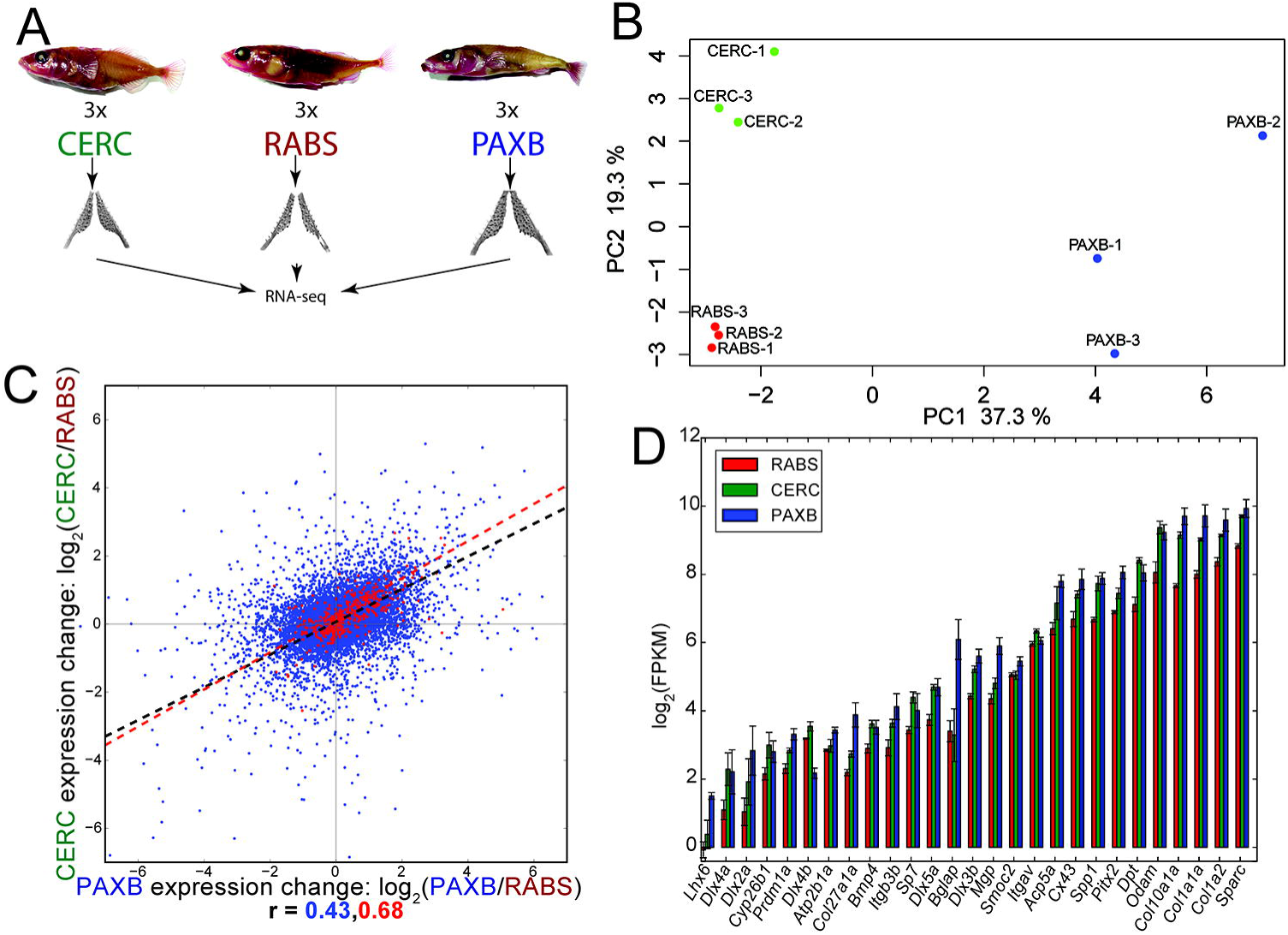
Evolved tooth gain in two freshwater populations. (A) Stickleback population locations. (B) Representative Alizarin red stained adult lab-reared sticklebacks (top, scale bars = 1 cm) and dissected ventral pharyngeal tooth plates (scale bars = 100μm). (C) Total ventral pharyngeal tooth number of 35–50 millimeter standard length lab-reared adult fish from each population.

To estimate the genomic relatedness of these populations, we resequenced the genomes of three marine and six freshwater sticklebacks from the four different populations (Table S2). We aligned the resulting reads to the stickleback reference genome [31] using Bowtie2 [42], and called variants using the Genome Analysis Toolkit (GATK) [43–45]. As it has been previously shown that Pacific marine stickleback populations are an outgroup to freshwater populations from Canada and California [31], we hypothesized the two high-toothed populations would be more related to each other genomically than either marine population. A maximum-likelihood phylogeny constructed using genome-wide variant data cleanly separated freshwater populations from each other and from marine fish (Fig S1A). Principal component analysis of the genome-wide variants revealed that the first principle component explains nearly half (41.4%) of the overall variance, and separates benthic sticklebacks from both creek and marine fish (Fig S1B). The second principal component separated both freshwater populations from marine populations. These results further support the model that populations of freshwater sticklebacks used a combination of shared and independent genetic changes [31,32] when evolving a set of similar morphological changes in response to a new environment.

### Convergent evolution of gene expression

As morphological changes are often the result of changes in gene expression patterns and levels, we sought to identify evolved changes in gene expression during tooth development at stages soon after the evolved differences emerge [41]. We quantified gene expression in ventral pharyngeal dental tissue in the two high-toothed freshwater and an Alaskan low-toothed marine population using RNA-seq (Fig 2A, Table S3-S4). Principal component (PC) analysis of the resulting gene expression matrix showed a clustering of gene expression by population, with the first PC separating benthic samples, and the second PC separating both benthic and creek samples from marine, similar to the PC analysis of the genome-wide variants (Fig 2B) [46].

**Fig 2.**
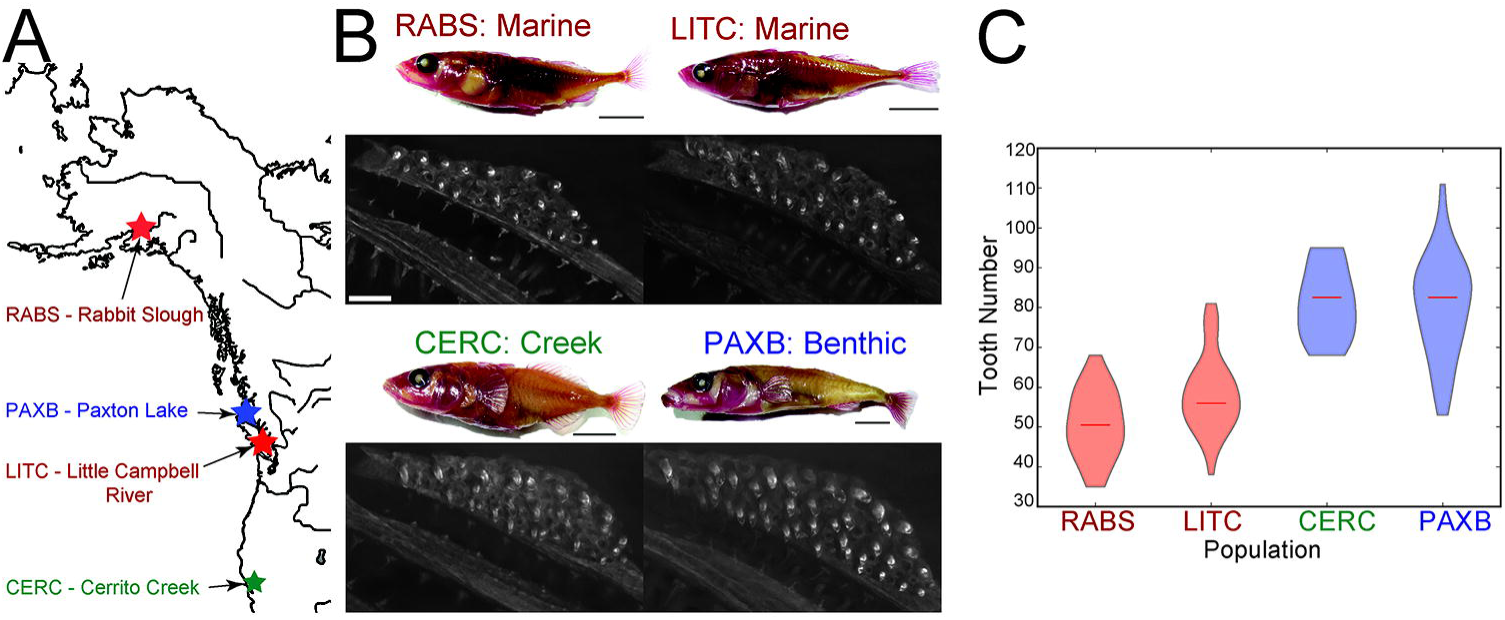
Convergent evolution of gene expression in dental tissue. (A) Ventral pharyngeal tooth plates from three different populations were dissected and gene expression quantified by RNA-seq. (B) Principal component analysis of dental tissue gene expression shows population specific expression profiles. (C) Freshwater dental tissue exhibited correlated gene expression changes for all genes (blue), with increased correlation observed for orthologs of genes known to be expressed during mammalian tooth development (red). (D) Expression of genes annotated as expressed in zebrafish teeth (zfin.org) which were significantly upregulated in one or both freshwater populations.

Given the convergently evolved morphological change of increases in tooth number, we hypothesized that convergent evolution has occurred at the gene expression level in freshwater dental tissue. To test this hypothesis, we compared the evolved change in gene expression in benthic dental tissue (benthic vs marine) to the evolved change in creek dental tissue (creek vs marine). At a genome-wide level, correlated changes in gene expression levels have evolved in the two high-toothed freshwater populations (Fig 2C, Spearman’s r = 0.43). We next asked if orthologs of genes implicated in tooth development in other vertebrates showed an increase in correlated evolved expression changes. We compared the gene expression changes of stickleback orthologs of genes in the BiteIt (http://bite-it.helsinki.fi/) [47] or ToothCODE (http://compbio.med.harvard.edu/ToothCODE/) [36] databases (hereafter referred to as the “BiteCode” gene set, Table S5), two databases of genes implicated in mammalian tooth development. Consistent with the conserved roles of gene regulatory networks regulating mammalian and fish teeth [48–51] and the major evolved increases in tooth number in both freshwater populations (Fig 1C), these predicted dental genes showed an increase in their correlated evolved gene expression change (Fig 2C red points, Spearman’s r = 0.68), and tended to have an overall increase in gene expression (Fig S2, *P* = 7.36e-6, GSEA, see methods). We also examined the expression levels of genes whose orthologs are annotated as being expressed in zebrafish pharyngeal teeth (www.zfin.org). Within this gene set, 27 of 40 genes were significantly more highly expressed in at least one freshwater population, with no genes expressed significantly higher (as determined by cuffdiff2 [52–55], see Materials and Methods) in marine samples than either freshwater population (Fig 2D).

### Increased freshwater expression of stem cell maintenance genes

Tooth development is controlled by several deeply conserved developmental signaling pathways [49,51]. To test whether expression changes in the components of specific developmental signaling pathways have evolved in the two high-toothed freshwater populations, we next analyzed the expression levels of stickleback orthologs of genes implicated in mammalian tooth development and annotated as components of different signaling pathways [36]. When comparing gene expression levels in freshwater dental tissue to marine dental tissue, genes annotated as part of the TGF-ß signaling pathway displayed significantly increased expression in freshwater dental tissue (Fig S3A-F).

Since these two freshwater populations have a largely different developmental genetic basis for their evolved tooth gain [30], we next asked whether any pathways were upregulated or downregulated specifically in one freshwater population. When comparing the expression of genes in benthic dental tissue to expression in creek or marine dental tissue, genes not only in the TGF-ß pathway, but also in the WNT signaling pathway, displayed significantly increased expression, consistent with the differing genetic basis of tooth gain in these populations (Fig S3B). Genes upregulated in freshwater dental tissue were enriched for Gene Ontology (GO) terms involved in anatomical structure development, signaling, and regulation of cell proliferation (Fig S4A, Table S6). Genes upregulated in benthic dental tissue over marine were enriched for GO terms involved in cell proliferation, division and cell cycle regulation, as well as DNA replication (Fig S4B, Table S7), while genes upregulated in creek over marine were enriched for GO terms involved in cell locomotion, movement, and response to lipids (Fig S4C, Table S8).

As teeth are constantly being replaced in polyphyodont adult fish, potentially due to the action of dental stem cells [40], we hypothesized that genes involved in stem cell maintenance have evolved increased expression in freshwater tooth plates, given the higher rate of newly forming teeth previously found in adults [30], and the possibly greater number of stem cell niches in high-toothed fish (Cleves et al 2018 under review, see Supplemental Data File 2). We further hypothesized that since teeth are developmentally homologous to hair, perhaps an ancient genetic circuit regulating vertebrate placode replacement controls both fish tooth and mammalian hair replacement. For example, the *Bmp6* gene, previously described as expressed in all stickleback teeth [41] was significantly upregulated in creek fish, consistent with the evolved major increases in tooth number in this population (Table S4). In contrast, no such significant upregulation was observed in the expression of benthic *Bmp6* (Table S4), consistent with the observed evolved *cis*-regulatory decrease in benthic *Bmp6* expression [41]. Further supporting this hypothesis, the expression of the stickleback orthologs of a previously published set of mouse hair follicle stem cell (HFSC) signature genes [56] were significantly upregulated in freshwater dental tissue (Fig S3A). Creek dental tissue displayed a small but significant increase in expression of this set of HFSC orthologs relative to both benthic and marine samples (Fig S3C).

In cichlid fish, pharmacology experiments revealed that reductions in tooth density can be accompanied by concomitant increases or decreases in taste bud density [39]. To begin to test whether derived high-toothed stickleback populations have also evolved significantly altered levels of known taste bud marker gene expression, we examined the expression levels of known taste bud markers *Calbindin2* and *Phospholipase Beta 2* [57], as well as taste receptors such as *Taste 1 Receptor Member 1, Taste 1 Receptor Member 3*, and *Polycystin 2 Like 1* [58]. Although four of these five genes had detectable significant expression changes between different populations, no consistent freshwater upregulation or downregulation of taste bud marker genes was seen (Fig S5).

### *Cis* and t*rans* regulatory changes in gene expression

Evolved changes in gene expression are due to a combination of *cis* acting changes that are linked to the genes they act on, and *trans* acting changes which usually are genetically unlinked to the gene or genes they regulate. Since the genetic basis of freshwater tooth gain mapped to non-overlapping intervals in these two populations [30] (Cleves et al 2018 under review, see Supplemental Data File 2), we hypothesized that the observed shared freshwater gene expression changes were the result of a similar *trans* environment, but a largely different set of *cis* changes. To test this hypothesis, we measured evolved *cis* expression changes in marine-freshwater F1 hybrids, which have marine and freshwater alleles present in the same *trans* environment. We raised both creek-marine and benthic-marine F1 hybrids to the late juvenile stage, dissected their ventral pharyngeal tooth plates, then generated and sequenced five barcoded RNA-seq libraries per population (10 total). We then quantified the *cis* expression change as the ratio of the number of reads mapping uniquely to the freshwater allele of a gene to the number of uniquely mapping marine reads (Fig 3A, Table S9-11). *Trans* expression changes were calculated by factoring the *cis* change out from the overall parental expression change [19].

**Fig 3.**
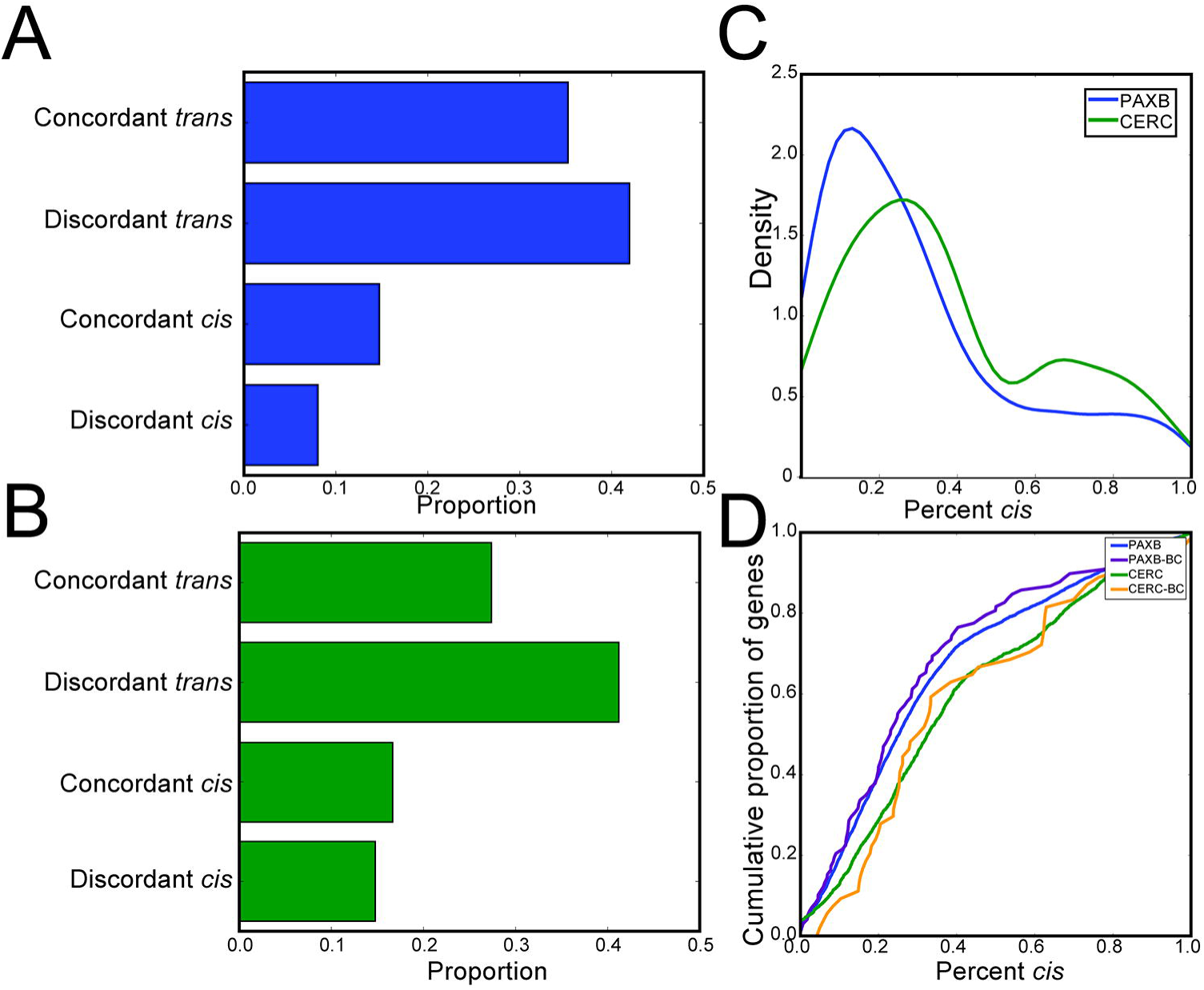
Evolved changes in *cis*-regulation. (A) Ventral pharyngeal tooth plates from marine-creek and marine-benthic lake F1 hybrids were dissected and *cis* regulatory changes assayed using phased RNA-seq reads. (B) Density plot showing the measured *cis*-regulatory changes. Neither population displayed a significant allelic bias, as measured by a Wilcoxon signed-rank test. (C-D) Gene expression changes in both parental and hybrid dental tissue – genes are color-coded based on the role of *cis* and/or *trans* change in benthic (C) or creek (D) dental tissue. Dashed line indicates the first principal component axis.

We found 11,832 and 8,990 genes in benthic and creek F1 hybrids, respectively, that had a fixed marine-freshwater sequence difference which had more than 20 total reads mapping to it. We observed no significant bias towards either the marine or freshwater allele in either set of F1 hybrids (Fig 3B). We next classified genes into one of four categories (*cis* change only, *trans* change only, concordant *cis* and *trans* changes, discordant *cis* and *trans* changes). We found 1640 and 1116 benthic (Fig. 3C) and creek (Fig. 3D) genes, respectively, with only significant *cis* changes, and 1873 and 1048 genes, respectively, with only significant *trans* changes. We also found 478 and 359 genes with significant *cis* and *trans* changes in the same direction, which we term concordant changes in gene expression. Conversely, we found 772 and 607 genes with significant *cis* and *trans* changes in opposing directions, which we termed discordant changes. Thus, overall, *trans* regulatory changes are more common than *cis* changes in the evolution of dental tissue gene expression in both freshwater populations. Additionally, discordant *cis* and *trans* changes were more common in both populations, suggesting selection for stable levels of gene expression.

### *Trans* regulatory changes dominate

We next wanted to determine the relative contribution of *cis* and *trans* gene expression changes to evolved changes in gene expression. We restricted our analysis to differentially expressed genes (as determined by cuffdiff2 [52]) to examine only genes with a significant evolved difference in gene expression and quantifiable (i.e. genes with transcripts containing a polymorphic variant covered by at least 20 reads) *cis* and *trans* expression changes. When evolving a change in gene expression, the *cis* and *trans* regulatory basis for this change can be concordant (*cis* and *trans* effects both increase or decrease expression) or discordant (*cis* effects increase and *trans* decrease or vice versa). We hypothesized that genes would tend to display more discordant expression changes, as stabilizing selection has been found to buffer gene expression levels [17,22,59]. To test this hypothesis, we binned genes into a 2x2 contingency table, with genes classified as *cis* or *trans* based on which effect controlled the majority of the evolved expression change, and discordant or concordant based on the direction of the *cis* and *trans* changes (Fig 4A, B). In the creek population, significantly more discordant changes than expected by a neutral mode*l* (*P* =1.35e-7, binomial test) have evolved. In both populations, we found increased discordant changes when the *trans* effect is larger than the *cis* effect (*P* =1.29e-7, 1.44e-13, benthic and creek respectively, binomial test). In both populations, we observe the opposite (an enrichment of concordant changes) when the *cis* effect is stronger, relative to the ratio when the *trans* effect is dominant (*P* =1.34e-36, 8.2e-11 benthic and creek respectively, binomial test). When considering all (not just differentially expressed) genes with quantifiable *cis* and *trans* expression changes, discordant changes dominated regardless of the relative strength of the *cis* effect (Fig S6).

**Fig 4.**
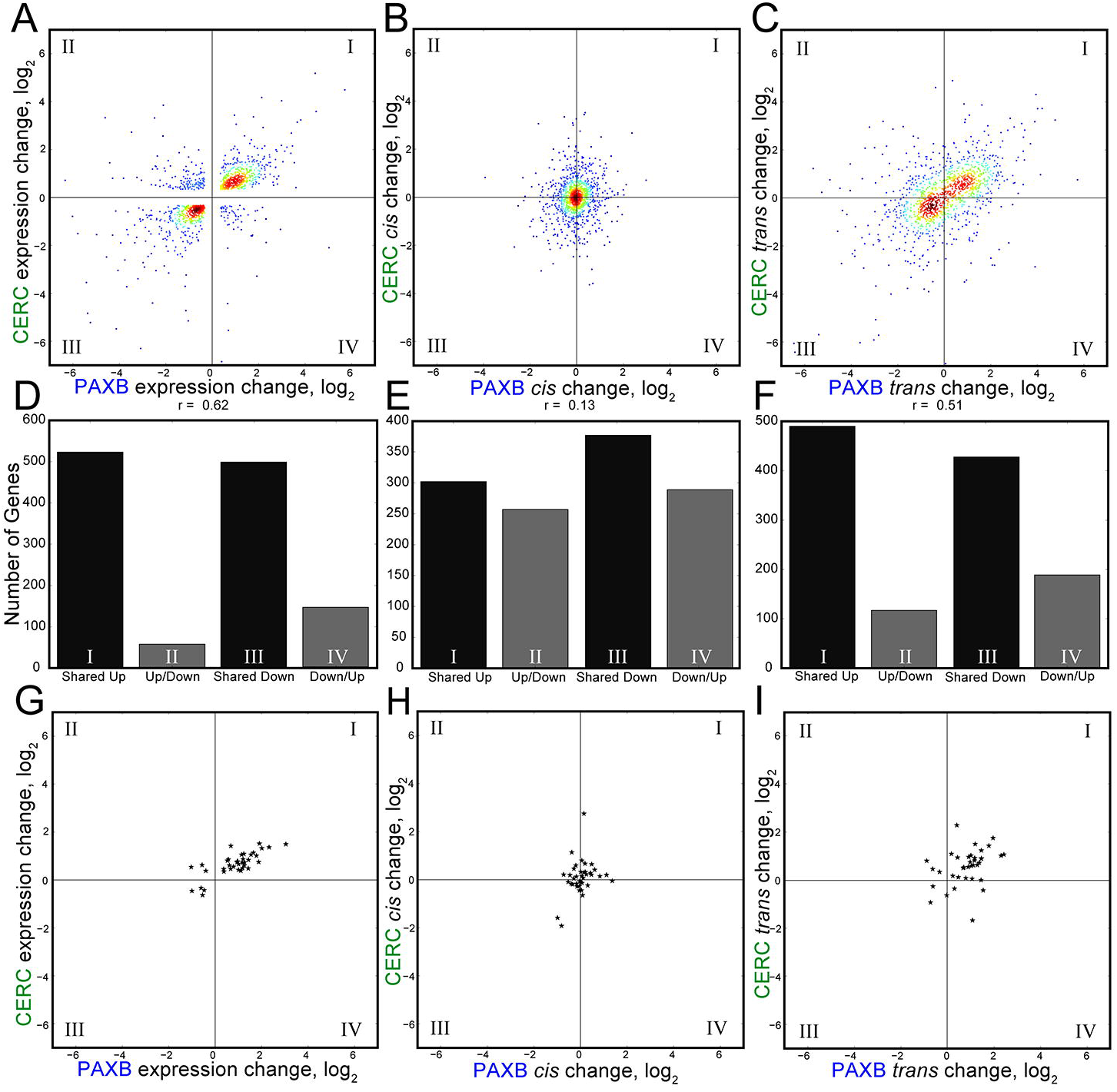
*Trans* changes predominate evolved dental gene expression changes. (A-B) Proportion of differentially expressed genes displaying opposing and concordant *cis* and *trans* changes in benthic (A) or creek (B) dental tissue. Genes whose expression differences were mostly explained by *cis* changes tended to be more concordant (*P* =5.0e-17, 0.002 for benthic and creek, respectively) than those mostly explained by *trans* changes. (C) Density of the relative percentage of gene expression differences which are explained by *cis* changes in benthic and creek dental tissue. (D) Cumulative percentage of percentage of gene expression due to *cis* changes. Genes in creek samples display a higher percentage *cis* change than genes in benthic samples (*P* = 1.25e-22, Mann-Whitney U test).

If all gene expression changes were due to changes only in *cis*, we would expect to see the measured *cis* ratios in the hybrids match the parental expression ratios. Instead, in both cases of evolved change, we saw parental expression ratios of a greater magnitude than F1 hybrid ratios, indicating a stronger contribution of *trans* changes to overall gene expression changes (Fig 3C-D). Indeed, when we examined the overall percentage of expression changes of differentially expressed genes that were due to changes in *cis*, we observed median per gene values of only 25.2% and 32.5% of benthic and creek gene expression changes, respectively (Fig 4C). Comparing the expression levels of orthologs of known dentally expressed genes from the BiteIt [47] and ToothCODE [36] databases revealed a similarly small number of gene expression changes explained by changes in *cis*, relative to the genome-wide average (Fig 4D). Evolved changes in creek gene expression were more due to changes in *cis* than benthic genes (Fig 4D, *P* = 1.25e-22, Mann-Whitney U test). Thus, *trans* effects on gene expression dominate the evolved freshwater gene expression changes.

### *Trans* regulatory changes are more likely to be shared between freshwater populations

We next wanted to test the hypothesis that the shared freshwater gene expression changes were primarily due to shared *trans* changes, rather than shared *cis* changes. We first compared the overall expression levels of genes called differentially expressed between benthic and marine as well as creek and marine. Similar to the genome-wide comparison, we found a highly significant non-parametric correlation coefficient (Spearman’s r = 0.62, *P* =1.2e-132) for the expression change of these shared differentially expressed genes (Fig 5A). When comparing the benthic *cis* changes to the creek *cis* changes, however, we found a much lower (though still significant) correlation coefficient (Spearman’s r = 0.13, *P* =5.1e-6) (Fig 5B). When comparing the calculated *trans* changes for these shared differentially expressed genes, we observed much higher correlation coefficient (Spearman’s r = 0.51, *P* =1.2e-80) (Fig 5C). When comparing all, not just differentially expressed, genes, *trans* changes are still likely to be more shared than *cis* (Fig S7). Additionally, 35/38 of the shared differentially expressed putative dental genes have shared regulatory increases or decreases in both freshwater populations relative to marine in overall expression difference, with 32/38 in *trans*, but only 25/38 in *cis* (Fig 5D-I). Thus, the *trans* effects on evolved gene expression are more likely to be shared by both freshwater populations than the *cis* changes.

**Fig 5.**
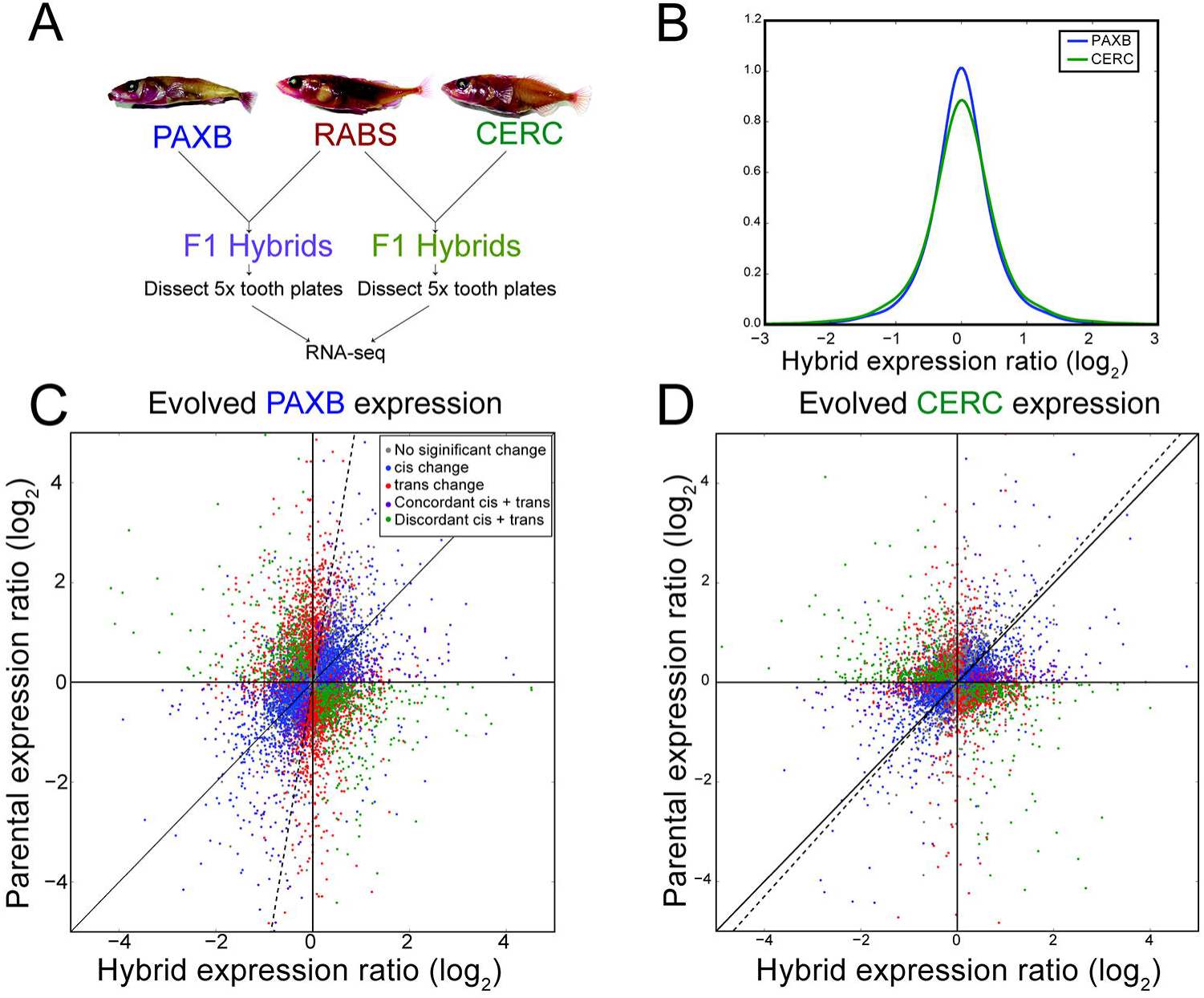
*Trans* changes are more likely to be shared across populations. (A) Genes with significantly different evolved expression in both freshwater populations relative to marine fish, showing significantly correlated changes in gene expression in benthic and creek dental tissue. (B) Freshwater dental tissue had a significant but small number of shared *cis*-regulatory changes. (C) Freshwater dental tissue showed significantly correlated changes in *trans* expression changes. A-C show genes with significant expression changes between populations and quantifiable (i.e. genes with transcripts containing a polymorphic SNP covered by at least 20 reads) *cis*-regulatory changes in both populations. Density (color) was estimated with a Gaussian kernal density estimator. BiteCode genes (see Methods) are indicated with black stars. D-F Bar graphs show the number of genes with shared or divergent expression patterns from the above panels. G-I are similar to A-C, but show only genes in the BiteCode gene set.

## Discussion

We sought to test the relative contribution of *cis* and *trans* gene regulatory changes during convergent evolution of tooth gain, as well as to ask whether the same or different regulatory changes underlie evolved changes in gene expression during this case of convergent evolution. We quantified the overall regulatory divergence, as well as the specific contribution of *cis* and *trans* changes, between ancestral low-toothed marine and two different independently derived populations of high-toothed freshwater sticklebacks. Similar overall changes in gene expression have evolved in both freshwater populations, especially in orthologs of known dental regulators in mammals. In this system, *trans*-regulatory changes play a larger role than *cis* changes in both populations. Furthermore, *trans* acting changes were much more likely to be shared between freshwater populations than *cis* changes, suggesting the two high-toothed populations evolved their similar gene expression patterns through independent genetic changes.

### Convergent evolution of dental gene expression

Convergent evolution at the gene expression level occurs when similar gene expression levels evolve in different populations. Both the creek and benthic stickleback populations have adapted from an ancestral marine form to their current freshwater environments. The genomic nature of their derived changes appears largely divergent, with major axis of variation separating benthic genomes from the geographically proximal marine populations (LITC), as well as the more distant marine (RABS) and creek populations. However, when looking at the gene expression basis of their convergently evolved gain in tooth number, orthologs of genes implicated in mammalian dental development showed strong correlated freshwater gains in expression. This correlation suggests both that sticklebacks deploy conserved genetic circuits regulating tooth formation during tooth replacement, but also that both populations have convergently evolved changes to similar downstream transcriptional circuits resulting in a gain of tooth number.

Though both freshwater populations showed strongly correlated changes in evolved gene expression at the *trans* regulatory level, the *cis* changes were largely not shared across populations. This was especially true for putative dentally expressed genes with evolved expression changes – the vast majority of the *trans* but not *cis* expression changes were shared between both freshwater populations. This suggests that the similar freshwater gene expression patterns evolved through independent genetic changes. It is possible that the small number of shared *cis* changes are sufficient to drive the observed changes to the overall *trans* regulatory environments. However previous work has shown that the genetic basis of tooth gain in these two populations is distinct [30] (Cleves et al 2018 under review, see Supplemental Data File 2), and it seems parsimonious that the genetic basis of a gain in dental gene expression is also mostly independent. Thus, convergent freshwater gene expression changes appear to be largely due to distinct, independent population-specific regulatory changes. This finding suggests that there are many regulatory alleles that are accessible during the evolution of an adaptive trait.

### *Trans* effects dominate

Other studies have used RNA-seq to compare the relative contribution of *cis* and *trans*-regulatory changes in the evolution of gene expression. In mice, evolved gene expression changes in the liver [18] and the retina [60] were driven primarily by *cis*-regulatory changes. In *Drosophila*, work on organismal-wide evolved gene expression changes on the genome-wide level has shown the opposite, with *trans*-regulatory effects playing a larger role in the evolution of gene expression [19,22]. Other studies have found *trans* effects contribute more to intraspecific comparisons, while *cis* effects contribute more to interspecific comparisons [61]. Consistent with this, we observe *trans* effects dominating in both of our intraspecific comparisons.

Another key distinction could be that *cis*-regulatory effects dominate when looking at more cellularly homogenous tissues, while *trans*-regulatory effects dominate when looking at more heterogeneous tissues. Stickleback tooth plates likely fall into an intermediate category, less heterogenous in cell type composition than a full adult fly or fly head, but more heterogeneous than a specialized tissue such as the mouse retina. Overall, freshwater tooth plates are more morphologically similar to each other than marine, with freshwater tooth plates possessing a larger area, increased tooth number, and decreased intertooth spacing [30,41]. Freshwater tooth plates likely have more similar cell type abundances and compositions (e.g. more developing tooth germs with inner and outer dental epithelia, and odontogenic mesenchyme) compared to each other than to marine tooth plates. Similar cell types tend to have similar gene expression patterns, even when compared across different species [62]. Much of the shared freshwater increase in dental gene expression could be due to an increase in dental cell types in both freshwater populations. As other evolved changes to stickleback morphology have been shown to be due to *cis* regulatory changes to key developmental regulatory genes [8,33,41,63], this *trans* regulatory increase in cell type abundance could be due to a small number of *cis* regulatory changes. These initially evolved developmental regulatory changes could result in similar downstream changes in the developmental landscape, resulting in the shared increase in dental cell types. Consistent with this interpretation, stickleback orthologs of genes known to be expressed during mammalian tooth development were found here to have a much greater incidence of convergently evolved increase in *trans* regulatory gene expression.

### Compensatory *cis* and *trans*

Previous studies [17,18] have shown compensatory *cis* and *trans* changes are essential for the evolution of gene expression. These findings are consistent with the idea that the main driving force in the evolution of gene expression is stabilizing selection [59] where compensatory changes to regulatory elements are selected for to maintain optimal gene expression levels. In both benthic and creek dental tissue, when considering all genes with a quantifiable (i.e. polymorphic and covered by ~20 reads, see Methods) *cis* effects, discordant compensatory *cis* and *trans* changes were far more common than concordant ones. This trend could be driven by some initial selection on pleiotropic *trans* changes, followed by selection for compensatory *cis* changes to restore optimal gene expression levels [17,18,22]. However, the *trans*, but not the *cis*, evolved changes in gene expression were highly shared among the two freshwater populations. Thus, collectively our data support a model where two independently derived populations have convergently evolved both similar genome-wide expression levels as well as ecologically relevant morphological changes through different genetic means.

### Potential parallels between teeth and hair regeneration

Creek and benthic sticklebacks have an increased rate of new tooth formation in adults relative to their marine ancestors [30]. In constantly replacing polyphyodonts, it has been proposed that teeth are replaced through a dental stem cell intermediate [37,38]. A strong candidate gene underlying a large effect benthic tooth quantitative trait locus (QTL) is the secreted ligand *Bone Morphogenetic Protein 6* (*Bmp6*) [41] (Cleves et al 2018 under review, see Supplemental Data File 2), which is also a key regulator of stem cells in the mouse hair follicle [56]. Freshwater dental tissue displayed significantly increased expression of known signature genes of mouse hair follicle stem cells, perhaps reflecting more stem cell niches supporting the higher tooth numbers in freshwater fish. Genes upregulated in freshwater dental tissue also were significantly enriched for GO terms involved in the cell cycle and cell proliferation. Together these findings suggest that both freshwater populations have evolved an increased tooth replacement rate through an increased activity or abundance of their dental stem cells, and also suggest the genetic circuitry regulating mammalian hair and fish tooth replacement might share an ancient, underlying core gene regulatory network.

## Materials and Methods

### Stickleback husbandry

Fish from all populations were raised in 110L aquaria in brackish water (3.5g/L Instant Ocean salt, 0.217mL/L 10% sodium bicarbonate) at 18°C in 8 hours of light per day. Young fry [standard length (SL) < 10 millimeters (mm)] were fed a diet of live *Artemia*, early juveniles (SL ~10 - 20 mm) a combination of live *Artemia* and frozen *Daphnia*, and older juveniles (SL > ~20 mm) and adults a combination of frozen bloodworms and *Mysis* shrimp. Experiments were approved by the Institutional Animal Care and Use Committee of the University of California-Berkeley (protocol # R330).

### Skeletal staining and imaging

Sticklebacks were fixed in 10% neutral buffered formalin overnight at 4°C. Fish were washed once with water and then stained in 1% KOH, 0.008% Alizarin Red for 24 hours. Following a water rinse, fish were cleared in 0.25% KOH, 50% glycerol for 2–3 weeks. Branchial skeletons were dissected as previously described [64]. Pharyngeal teeth were quantified with fluorescent illumination using a TX2 filter on a Leica DM2500 microscope. Representative tooth plates were created using montage z-stacks on a Leica M165 FC using the RhodB filter. Adult fish were imaged using a Canon Powershot S95. Some tooth count data from the CERC, RABS, and PAXB populations; n = 11, 13, 29, respectively, (see Table S1) have been previously published [30].

### DNA preparation and genome resequencing

Caudal fin tissue was placed into 600µl tail digestion buffer [10mM Tris pH 8.0, 100mM NaCl, 10mM EDTA, 0.05% SDS, 2.5µl ProK (Ambion AM2546)] for 12 hours at 55°C. Following addition of 600 µl of 1:1 phenol:chloroform solution and an aqueous extraction, DNA was precipitated with the addition of 1ml 100% ethanol, centrifuged, washed with 75% ethanol, and resuspended in water. 50ng of purified genomic DNA was used as input for the Nextera Library prep kit (Illumina FC-121-1031), and barcoded libraries were constructed following the manufacturer’s instructions. Library quality was verified using an Agilent Bioanalyzer. Libraries were pooled and sequenced on an Illumina HiSeq 2000 (see Table S2 for details).

### RNA purification and creation of RNA-seq libraries

Late juvenile stage female sticklebacks (SL ~40mm) were euthanized in 0.04% Tricaine. Dissected [64] bilateral ventral pharyngeal tooth plates were placed into 500µl TRI reagent, then incubated at room temperature for 5 minutes. Following addition of 100µl of chloroform, a further 10 minute incubation and centrifugation, the aqueous layer was extracted. Following addition of 250µl isopropyl alcohol and 10 minute incubation, RNA was precipitated by centrifugation, washed with 75% EtOH, and dissolved in 30ul of DEPC-treated water. RNA integrity was assayed by an Agilent Bioanalyzer. 500ng of RNA from each fish was used as input to the Illumina stranded TruSeq polyA RNA kit (Illumina RS-122-2001), and libraries were constructed following the manufacturer’s instructions. Library quality was analyzed on an Agilent Bioanalyzer, and libraries were pooled and sequenced on an Illumina HiSeq2000 (see Table S3).

### Gene expression quantification and analysis

RNA-seq reads were mapped to the stickleback reference genome [31] using the STAR aligner [65] (version 2.3, parameters = --alignIntronMax 100000 --alignMatesGapMax 200000 --outFilterMultimapNmax 20 --outFilterMismatchNmax 999 --outFilterMismatchNoverLmax 0.04 --outFilterType BySJout), using ENSEMBL genes release 85 as a reference transcriptome. The resulting SAM files were sorted and indexed using Samtools version 0.1.18 [66], PCR duplicates were removed, read groups added and mate pair information fixed using Picard tools (version 1.51) (http://broadinstitute.github.io/picard/) with default settings. Gene expression was quantified with the Cufflinks suite (v 2.2.1) [52–55] using ENSEMBL genes as a reference transcriptome, with gene expression quantified with cuffquant (-u --library-type fr-firststrand) and normalized with cuffnorm. Differentially expressed genes were found using cuffdiff2, with parameters (-u --FDR.1 --library-type fr-firststrand, using the reference genome for bias correction). Genes with a mean expression less than 0.1 FPKM were filtered from further analysis.

### Gene set and gene ontology enrichment

The BiteCode gene set was generated by combining all genes in the BiteIt (http://bite-it.helsinki.fi/) or ToothCODE (http://compbio.med.harvard.edu/ToothCODE/) [36] databases. Stickleback orthologs or co-orthologs were found using the annotated names of ENSEMBL stickleback genes. Gene set expression change statistical enrichment was done as previously described [67]. Briefly, a t-test was performed for each gene to test for a difference in mean expression between the two treatments. The resulting t-values were subject to a 1-sample t-test, with the null model that the mean of the t-values was 0. Cutoffs were validated using 10,000 bootstrapped replicate gene sets drawn from the same gene expression matrix. Stickleback orthologs of mouse or human genes were determined using annotated ENSEMBL orthologs. Sorted lists of genes, ranked by log_2_ expression change in benthic dental tissue relative to marine, creek relative to marine, or the mean of creek and benthic relative to marine, were generated using the measured gene expression data. Gene Ontology enrichment was done using Gorilla [68,69], and results were visualized using REVIGO [70].

### Detection of genomic and transcriptomic variants

Genomic resequencing reads were aligned to the stickleback reference genome [31] using the bwa aln and bwa sampe modules of the Burrows-Wheeler Alignment tool (v 0.6.0-r85) [71]. Resulting SAM files were converted to BAM files, sorted and indexed by Samtools version 0.1.18 [66], with PCR duplicates removed by Picard tools. GATK’s (v3.2–2) IndelRealigner (parameter: ‘-LOD 0.4’), BaseRecalibrator, and PrintReads were used on the resulting BAM files. BAM files from the above RNA-seq alignment were readied for genotype calling using GATK’s SplitNCigarReads, BaseRecalibrator, and PrintReads. Finally, the UnifiedGenotyper was used to call variants from the RNA-seq and DNA-seq BAM files, with parameters (-stand_call_conf 30 -stand_emit_conf 30 -U ALLOW_N_CIGAR_READS --genotype_likelihoods_model BOTH) [43,45].

Following final variant calling and detection, pseudo-transcriptomes were created for each F1 hybrid. The pseudo-transcriptomes consist of the predicted sequence for each allele within an F1 hybrid, with all predicted splicing variants of a gene collapsed to a single transcript. A variant was added to the pseudo-transcriptome if and only if it was homozygous in the sequenced parents (or parent’s sibling in the case of the Alaskan marine parent of the Cerrito creek x Alaskan marine F1 hybrids) and called heterozygous in the F1 hybrid.

### *Cis* and *trans* regulatory divergence quantification

RNA-seq reads from F1 hybrid sticklebacks were aligned to the individual’s pseudo-transcriptome using STAR (v 2.3) with the parameters: --outFilterMultimapNmax 1 and --outFilterMultimapScoreRange 1. By only looking at uniquely aligning reads, we ensured we only considered reads which overlapped a heterozygous variant site. Counting these unique reads minimizes double counting a single read that supports two different variant positions. Total *cis* divergence in each F1 hybrid was quantified by comparing the number of reads mapping uniquely to each allele in the pseudo-transcriptome.

Following *cis* divergence quantification in all F1 hybrids, we considered the overall *cis* change in the different freshwater populations. Genes which only had 20 or fewer uniquely mapping reads across all replicates were filtered from further analysis. We excluded genes with more than a 32-fold change, as a manual inspection revealed these to be either genotyping errors or mitochondrial genes. Reported *cis* ratios were calculated by comparing the ratio of uniquely mapped freshwater reads to uniquely mapped marine reads. Evolved *trans* changes were quantified as the difference between the log of the overall gene expression change between the freshwater and marine parents and the log of measured *cis* freshwater expression change. Percent *cis* change was calculated as the absolute value of the log of the *cis* change divided by the sum of the absolute value of the log of the *cis* change and the absolute value of the log of the *trans* change. Statistical significance of *cis* changes was determined by a binomial test comparing overall reads mapping to the freshwater allele to a null model of no *cis* divergence, with a false discovery rate of 1% applied using the Benjamini-Hochberg method. Statistical significance of *trans* changes was determined by a G-test, comparing the expected (based on the measured *cis* change) and observed ratios of marine and freshwater, with a 1% false discovery rate.

## Data Availability

All sequencing reads are available on the Sequence Read Archive (XXXXXX). All scripts used for analysis are available on GitHub (xxxxx).

## Acknowledgements

We thank Nikunj Donde for collecting a subset of tooth count data.

---

**Fig S1. Independent freshwater evolutionary history.** (A) Genome-wide maximum-likelihood phylogeny created from genomic resequencing data. Wild-caught fish are non-italicized. All nodes have 100% bootstrapping support. (B) Principal component analysis of genome-wide genotypes separates marine and creek populations from the benthic lake population, with the 2^nd^ PC separating marine and freshwater populations.

**Fig S2. Freshwater upregulation of putative dental gene**s. (A) Benthic upregulation of BiteCode genes (*P* = 9.8e-3, GSEA). (B) Creek upregulation of BiteCode genes (*P* = 2.1e-5, GSEA). (C) Benthic and creek upregulation of BiteCode genes (*P* = 5.1e-6, GSEA).

**Fig S3. Concerted changes in stem cell markers and signaling pathway**s. (A-F) Changes in gene expression changes of genes annotated as components of the indicated signaling pathways (BMP, FGF, SHH, WNT, ACT, TGFB, NOTCH, or EDA) [36] or orthologs of a described set of mouse hair follicle stem cell signature genes (HFSC) [56]. Violin plots show the mean expression change of genes in the pathway. (A) Change in freshwater (benthic + creek) relative to marine. (B) Benthic specific changes (benthic relative to creek + marine). (C) Creek specific changes (creek relative to benthic + marine). (D) Benthic evolved changes (benthic relative to marine) (E) Creek evolved changes (Creek relative to marine) (F) Benthic vs creek changes (benthic relative to creek).

**Fig S4. Gene ontology of freshwater upregulated genes.** (A-C) GO enrichment of genes upregulated in benthic (A), creek (B), or both (C). GO analysis was preformed using Gorilla [68], with the results visualized with Revigo [70].

**Fig S5. Expression of taste bud marker genes.** Expression levels of known taste bud marker genes in marine, benthic and creek tooth plates as assayed by RNA-seq. * indicates differentially expressed genes. Error bars are standard error of the mean.

**Fig S6. Compensatory changes dominate genes with no significant evolved gene expression difference.** (A-B) Proportion of genes with quantifiable (i.e. genes with transcripts containing a polymorphic SNP covered by at least 20 reads) hybrid expression displaying opposing and concordant *cis* and *trans* changes in benthic (A) or creek (B) dental tissue. Similar to Fig. 5, but here showing all genes, not just genes with significantly different expression levels compared to marine. *Trans* regulatory changes predominate, as do opposing over concordant changes. (C) Density plot of the percentage of gene expression changes explained by *cis*-regulatory changes.

**Fig S7. *Trans* changes are more likely to be shared across populations.** (A) Expression changes of genes with quantifiable (i.e. genes with transcripts containing a polymorphic SNP covered by at least 20 reads) hybrid expression in both freshwater populations relative to marine fish, showing significantly correlated changes in gene expression in benthic and creek tooth plates. (B) *cis* regulatory changes of genes with quantifiable hybrid expression expression in freshwater dental tissue overall do not display correlated evolved changes. (C) *trans* regulatory changes of genes with quantifiable hybrid expression in freshwater dental tissue. Density (color) was estimated with a Gaussian kernal density estimator. (D-F) Bar graphs show the number of genes with shared or divergent expression patterns from A-C. G-I are similar to A-C, but show only genes in the BiteCode gene set, revealing that these orthologs have evolved highly convergent changes in the two freshwater populations (G), despite non-convergent *cis* regulatory changes (H).

**Table S1. Population ventral pharyngeal tooth counts**

For each fish, the population, ecotype (freshwater or marine), total ventral pharyngeal tooth number (TVTP), total length (TL), standard length (SL), and whether data has been published [30] is shown.

**Table S2. Genomic DNA sequencing reads**

For each fish, population and biological replicate number (Fish), the total number of barcoded reads from each fish (reads), and number of reads that mapped and passed all filters (final mapped) is listed.

**Table S3. RNA-seq reads**

For each fish, population of parents and biological replicate number (sample), standard length (SL), total reads (generated by HiSeq2000 over two different runs (run1 and run2)), mapped reads (reads that mapped to the genome), and final reads (excludes reads filtered due to low quality or PCR duplication) is listed.

**Table S4. Overall gene expression in tooth plate**

Estimated abundance in in fragments per kilobases per million reads (FPKM) of ENSEMBL genes (rows) in ventral pharyngeal dental tissue from three individual fish from three populations (in columns). Mean expression (in FPKM) is shown after the 3 replicates. Log_2_(Pop1/Pop2) shows the fold-change in log_2_ of the estimated mean expression between the two populations. IsSig(Pop1/Pop2) indicates whether the difference was significant as reported by cuffdiff2.

**Table S5. BiteCode genes in sticklebacks**

A list of stickleback orthologs in the BiteIt [47] (http://bite-it.helsinki.fi/) or ToothCODE (http://compbio.med.harvard.edu/ToothCODE/) [36] databases.

**Table S6. GO process upregulated in freshwa**ter

Gene Ontology (GO) term category and name are given in GO term and description, with the p-value, q-value, and relative enrichment within genes upregulated in freshwater dental tissue reported by GOrilla [68].

**Table S7. GO process upregulated in benthic**

Gene Ontology (GO) term category and name are given in GO term and description, with the p-value, q-value, and relative enrichment within genes upregulated in benthic dental tissue reported by GOrilla [68].

**Table S8. GO process upregulated in creek**

Gene Ontology (GO) term category and name are given in GO term and description, with the p-value, q-value, and relative enrichment within genes upregulated in creek dental tissue reported by GOrilla [68].

**Table S9. F1 hybrid RNA-seq reads**

For each ventral pharyngeal tooth plate (VTP), population of parents and biological replicate number (sample), standard length (SL), total reads (generated by HiSeq2000), mapped reads (reads that mapped to the genome), final reads (excludes reads filtered due to low quality or PCR duplication), and unique reads (reads that mapped uniquely to one haplotype) is listed.

**Table S10. Benthic vs marine *cis* divergence**

Estimated gene expression change in *cis* in log_2_, benthic vs marine. Name is the reported ENSEMBL gene name. Log_2_(F/M) is the log_2_ of the ratio of freshwater vs marine reads mapping uniquely to the gene.

**Table S11. Creek vs marine *cis* divergence**

Estimated gene expression change in *cis* in log_2_, creek vs marine. Name is the reported ENSEMBL gene name. Log_2_(F/M) is the log_2_ of the ratio of freshwater vs marine reads mapping uniquely to the gene.

